# Inferring demography and selection in organisms characterized by skewed offspring distributions

**DOI:** 10.1101/440404

**Authors:** Andrew M. Sackman, Rebecca Harris, Jeffrey D. Jensen

## Abstract

The recent increase in time-series population genomic data from experimental, natural, and ancient populations has been accompanied by a promising growth in methodologies for inferring demographic and selective parameters from such data. However, these methods have largely presumed that the populations of interest are well-described by the Kingman coalescent. In reality, many groups of organisms, including viruses, marine organisms, and some plants, protists, and fungi, typified by high variance in progeny number, may be best characterized by multiple-merger coalescent models. Estimation of population genetic parameters under Wright-Fisher assumptions for these organisms may thus be prone to serious mis-inference. We propose a novel method for the joint inference of demography and selection under the Ψ-coalescent model, termed Multiple-Merger Coalescent Approximate Bayesian Computation, or MMC-ABC. We first quantify mis-inference under the Kingman and then demonstrate the superior performance of MMC-ABC under conditions of skewed offspring distribution. In order to highlight the utility of this approach, we re-analyzed previously published drug-selection lines of influenza A virus. We jointly inferred the extent of progeny-skew inherent to viral replication and identified putative drug-resistance mutations.

## Introduction

Elucidation of the underlying processes of evolution through the measurement of temporal changes in allele frequencies has remained a major focus of population genetics since the founding of the field (Fisher, 1930; Wright, 1931). Advancements in sequencing technologies over the last decade have dramatically increased the availability of genome-wide time-sampled polymorphism data for a wide variety of organisms, and several methods have been developed to analyze such data (Malaspinas et al., 2012; Mathieson and McVean, 2013; Foll et al., 2014a; Lacerda and Seoighe, 2014; Steinrücken et al., 2014; Ferrer-Admetlla et al., 2016; Schraiber et al., 2016; Shim et al., 2016; Rousseau et al., 2017). Of primary interest is the estimation of site-specific selection coefficients, and methods have been created that account for non-equilibrium demography and environmental fluctuations by, for example, accounting for effective population size, population structure, and changing selection intensities.

Time-series polymorphism data is generally available from three sources: experimentally evolved populations, clinical patient samples, and ancient specimens. Viruses are well-represented amongst such data, both owing to their obvious clinical relevance, as well as their short generation times, small genomes, and relatively high mutation rates. However, aspects of viral biology render the application of standard population genetic inference methods problematic. Namely, existing methodologies for analyzing time-sampled polymorphism data are generally developed around the Kingman coalescent framework and the Wright-Fisher (WF) model (Wright, 1931; Kingman, 1982) and are of highly questionable applicability to organisms typified by large variances in offspring distributions, or so-called “sweepstakes reproduction,” including not only viruses but many classes of prokaryotes, fungi, plants, and animals (reviewed in Tellier and Lemaire, 2014; Irwin et al., 2016).

In particular, the WF model assumes constant population size, random mating, non-overlapping generations, and Poisson offspring distributions with equal mean and variance. The Kingman coalescent is derived in the limit of the WF model and shares its assumptions. Reassuringly, population genetic statistics and methods developed under the Kingman have been shown to be robust to violations of WF assumptions (Möhle, 1998, 1999), and have been extended to incorporate selection, migration, and population structure (Neuhauser and Krone, 1997; Nordborg, 1997; Wilkinson-Herbots, 1998). However, large variance in offspring number (Eldon and Wakeley, 2006; Matuszewski et al., 2018), strong selection (Neher and Hallatschek, 2013; Schweinsberg, 2017), large sample sizes (Wakeley and Takahashi, 2003; Bhaskar et al., 2014), and recurrent selective sweeps (Durrett and Schweinsberg, 2004, 2005) may violate the critical assumption underlying the Kingman coalescent that only two lineages may coalesce at a time. Such a violation may produce genealogies that are characterized by multiple-lineage mergers. Thus, the analysis of genomic data from organisms characterized by highly skewed offspring distributions—such as viruses—may be prone to serious mis-inference if examined with traditional WF and Kingman based approaches, even under neutrality. In particular, the neutral multiple merger events induced by the reproductive biology of these organisms may be mistaken for multiple merger events induced by positive selection.

Though not widely utilized for inference, an alternative class of multiple-merger coalescent (MMC) models have been developed that are more general than the Kingman (*e.g.* Bolthausen and Sznitman, 1998; Pitman, 1999; Sagitov, 1999; Schweinsberg, 2000; Möhle and Sagitov, 2001), many being derived from Moran models generalized to allow multiple offspring per individual. Many of the recently derived MMC models form specific sub-classes of the Λ-coalescent, of which the Kingman is also a specific case in which only two lineages are allowed to merge in a generation (Donnelly and Kurtz, 1999; Pitman, 1999; Sagitov, 1999). It has been demonstrated that expectations under MMC models differ from those of the Kingman coalescent in several significant ways: effective population size (*N_e_*) does not scale linearly with census size (*N*) as it does under the Kingman (Huillet and Möhle, 2011); the site frequency spectrum (SFS) is skewed toward an excess of low-and high-frequency variants relative to the standard WF expectations, even under equilibrium neutrality (Eldon and Wakeley, 2006; Blath et al., 2016); and the fixation probability of new beneficial mutations approaches one as population size increases (Der et al., 2011).

Eldon and Wakeley (2006, 2008, 2009) introduced a specific case of the broader class of Λ MMC models, the Ψ-coalescent, under which the parameter *Ψ* describes the proportion of offspring in the population originating from a single parent in the previous generation. The Ψ-coalescent has been used in several instances to infer the strength and frequency of sweepstake events in marine organisms typified by Type-III survivorship curves (Eldon and Wakeley, 2006; Birkner et al., 2013; Blath et al., 2016; Matuszewski et al., 2018), and the expected SFS has been determined under both standard and non-equilibrium demography (Matuszewski et al., 2018).

Thus, we here introduce a novel statistical inference approach, termed Multiple-Merger Coalescent Approximate Bayesian Computation (MMC-ABC), for inferring population genetic parameters from time-sampled polymorphism data in populations subject to sweepstakes reproduction. MMC-ABC first characterizes the neutral demography of the population by generating genome-wide estimates of *N* and *Ψ*. It then estimates site-specific selection coefficients under the inferred sweepstakes model. We demonstrate that failing to account for skewed offspring distributions results in strong mis-inference of both demography and selection, and that MMC-ABC is capable of accurate joint estimation of offspring skew and selection coefficients even when the population size is not precisely known.

## Methods

### *N*_e_-based ABC method

The data *X* consist of allele frequency trajectories measured at *L* loci: *x_i_* (*i* = 1,…, *L*). The *N_e_*-based ABC methodology (modified from the method of Foll et al. (2014a)) infers genome-wide values of *N* and *Ψ* and *L* locus-specific selection coefficients *s_i_*(*i* = 1,…, *L*). At a particular locus *i*, we can approximate the joint posterior distribution as:

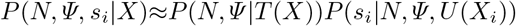

where *T* (*X*) = *T* (*X*_1_,…, *X_L_*) denotes summary statistics chosen to be informative about *N* and *Ψ* that are a function of all loci, and *U* (*X_i_*) denotes locus-specific summary statistics chosen to be informative about *s_i_*. A two-step ABC algorithm as proposed by Bazin et al. (2010) is used to approximate this posterior:

Step 1. Obtain an approximation of the density

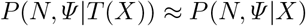

a. Simulate *L* trajectories for *J* populations *X_i,j_* using the starting frequencies from the first time point in each trajectory *x_i_* with *N* and *Ψ* for each trajectory randomly sampled from their priors and *J* equal to the total number of simulation replicates.
b. Compute *T* (*X_i,j_*) for each simulated population.
c. Retain the simulations with the smallest Euclidian distance between *T* (*X*) and *T* (*x*) to obtain a sample from an approximation to *P* (*N, Ψ |T* (*X*)) ≈ *P* (*N, Ψ |X*).
Step 2. For loci *i* = 1 to *i = L*:

a. Simulate *K* trajectories *X_i,k_* from a Ψ-coalescent model with *s_i_* randomly sampled from its prior and *N* and *Ψ* from the joint density obtained in step 1.
b. Compute *U* (*X_i,k_*) for each simulated trajectory.
c. Retain the simulations with the smallest Euclidian distance between *U* (*X_i_*) and *U* (*x_i_*) to obtain a sample from an approximation to *P* (*s_i_|N, Ψ, X_i_*)*P* (*N, Ψ |X*) = *P* (*N, Ψ, s_i_|X*).

As in the WF-ABC methodology of Foll et al. (2014b), we define *T* (*X*) as a single statistic, *Fs′*, an unbiased estimator of *N_e_*, given by Jorde and Ryman (2007):

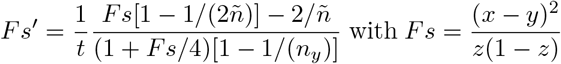

where *x* and *y* are the minor allele frequencies at the two time points separated by *t* generations, *z* = (*x* + *y*)/2, and *ñ* is the harmonic mean of the sample sizes *n_x_* and *n_y_* at the two time points expressed in the number of chromosomes (twice the number of individuals for diploids). We averaged *Fs′* values over sites and times to obtain a genome-wide estimator of *N_e_* = 1*/F s′* for haploids and *N_e_* = 1/2*Fs′* for diploids (Jorde and Ryman, 2007). Note that we use the common notation where *N_e_* corresponds to the effective number of individuals, and the corresponding number of chromosomes for diploids is 2*N_e_*.

In the second step of MMC-ABC, simulations are performed under a Ψ-coalescent model in SLiM version 3 (Haller and Messer, 2018, discussed in further detail in the next section) with an initial allele frequency and sample size matching those observed and with *N* and *Ψ* drawn from a joint posterior derived during Step 1. At each site we utilize two summary statistics derived from 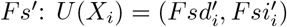 with *Fsd′* and *Fsi′* calculated, respectively, between pairs of time points where the allele considered is decreasing and increasing in frequency, such that at a given site *Fs′* = *Fsd′* + *Fsi′*. For the diploid model, we define the relative fitness as *w_AA_* = 1 + *s*, *w_Aa_* = 1 + *sh* and *w_aa_* = 1 where *h* denotes the dominance ratio (1 = dominant, 0.5 = codominance, 0 = recessive), and as *w_A_* = 1 + *s* and *w_a_* = 1 for the haploid model (Ewens, 2004).

### Forward simulation of populations under the Ψ-coalescent

Eldon and Wakeley (2006) described a model, the Ψ-coalescent, where each reproductive event in a population of size *N* is either, with probability 1 − *ε*, a standard WF event yielding a single offspring, or, with probability *ε*, a multiple-merger event yielding *ΨN* offspring. The probability *ε* = 1*/N^γ^* such that the coalescent history of a sample is dominated by multiple-merger events when 0 < *γ* < 2, and *γ* ≥ 2 produces a coalescent history typical of the Kingman. The rate at which *k* out of *n* lineages merge under the Ψ-coalescent is therefore (Tellier and Lemaire, 2014):

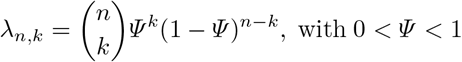

Under this model, *Ψ* has a straightforward biological interpretation. Namely, it is equal to the proportion of individuals in generation *t_i_* who are the offspring of a single individual in *t*_*i*−1_ (Eldon and Wakeley, 2006). We simulated populations evolving under a Ψ-coalescent model with SLiM. To circumvent the WF framework of SLiM, we utilized a system of subpopulations with migration to achieve the same effect as sweepstakes reproduction events. Each generation consists of three steps:

1. One individual is chosen from the population (A) and placed in a separate subpopulation (B) of size *N* = 1. The unidirectional migration rate from B to A is set to *Ψ*.
2. One WF generation occurs, with migration from subpopulation B resulting in a the chosen individual contributing *NΨ* of the individuals of the next generation of A. A series of mate choice callbacks within SLiM force the migration rate to be exact, rather than stochastic (see source code in the Supplementary Materials).
3. Subpopulation B is removed, and the next generation begins.

In Step 1 of MMC-ABC, a population of size *N* and skew *Ψ* (chosen from their prior distributions) is evolved with mutations reflecting the empirical data. The starting frequencies *x*_*i*,1_(*i* = 1,…, *L*) are identical to those observed during the first sampled time point. The frequency of each allele under consideration is output at each generation of the trajectories in *X*. In Step 2 of MMC-ABC, a population of size *N* and *Ψ* chosen from the joint posterior generated in part one is simulated with a single mutation of selection coefficient *s* chosen from its prior and starting at its observed initial frequency. The allele frequency is output each generation.

### Simulated data sets for testing performance of MMC-ABC

The data used for testing the performance of MMC-ABC were generated in one of two ways:

1. A diploid population of size *N* was first evolved under standard, neutral WF conditions for a burn-in period of 50,000 generations, and then evolved for a period of time under sweepstakes conditions. The frequencies of every segregating allele were output at the onset of sweepstakes conditions and at predetermined intervals for a set number of generations, including mutations present at the start of output as well as mutations that arose or fixed during the output period. Trajectories meeting minimum criteria (at least three informative time points, at least two consecutive time points with frequency greater than 0.01, and at least one time point with frequency higher than 0.025) were retained. Data were generated in this manner for testing the performance of Step 1 of MMC-ABC (joint estimation of *N* and *Ψ*). Unfiltered single-time point population data were used to generate the observed SFS data in Fig. 1.
2. Individual trajectories of mutations of a given starting frequency with selection coefficient *s* were modeled in a diploid population of size *N* with free recombination so that all sites were unlinked, with allele frequency trajectories and sweepstakes dynamics beginning in generation one. Trajectories generated in this manner were pooled into larger data sets for use in testing the performance of Step 2 of MMC-ABC (estimation of site-specific selection coefficients), with all allele trajectories beginning at a minor allele frequency of 10%—a low enough frequency that most neutral mutations should not fix, but high enough to ensure the availability of multiple informative time points for most trajectories.

**Figure 1.**
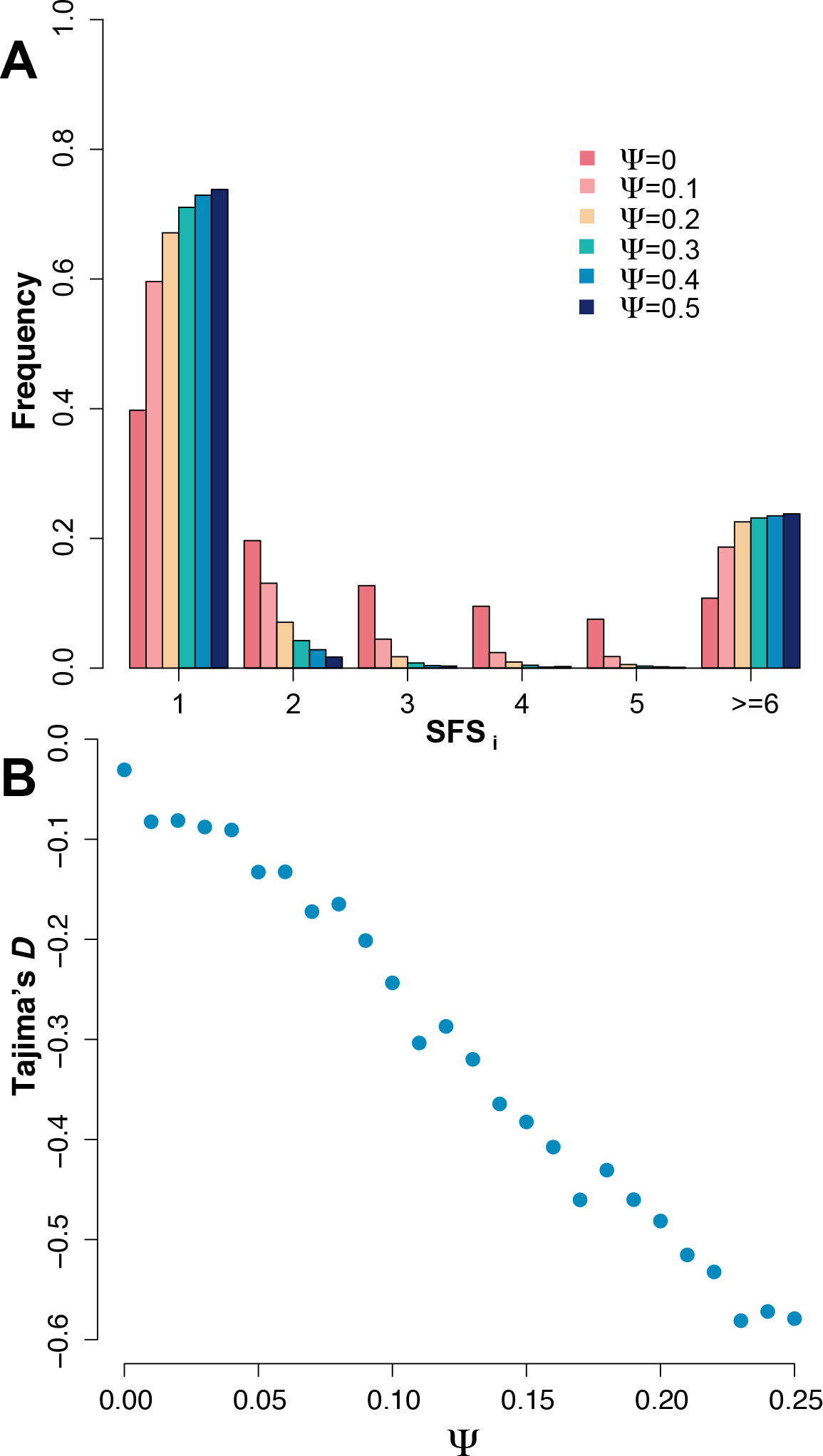
**A:** The site frequency spectrum (SFS) for Ψ ∈ {0; 0:1; 0:2; 0:3; 0:4; 0:5}, derived from values averaged over 100 replicate simulations at *N* = 1000 with sample size *n* = 250. **B:** The value of Tajima’s D for 0 ≤ Ψ ≤ 0:25 averaged across 100 replicate simulated populations with sample size *n* = 30. As shown, offspring skew strongly biases commonly used summary statistics, even under equilibrium neutrality.

We used a *Ψ* = 0.1 throughout our study, as this is close to the value estimated for experimentally evolved lines of influenza analyzed below. Additionally, at this level of skew, multiple mergers should dominate the coalescent history of a population without entirely eliminating all segregating variation.

### Analysis of drug-resistance in influenza A virus

We applied MMC-ABC to time-series polymorphism data from experimentally evolved populations of influenza A virus, originally described by Foll et al. (2014a). The data consist of population genomic sequencing from two control lineages and two lineages exposed to exponentially increasing concentrations of the influenza drug oseltamivir, reared on Madin-Darby canine kidney (MDCK) cells and sampled every thirteen generations. The data were previously analyzed with WF-ABC and putative drug-resistance mutations were identified. We reanalyzed the data with MMC-ABC for comparison.

### Data Availability

The source code and manual for MMC-ABC, along with the SLiM and python scripts used to generate our simulated data, will be made freely available upon acceptance from the “software” page of the Jensen Lab website: http://jjensenlab.org/software/. The raw data from the experimentally evolved influenza virus populations can be found at the ALiVE repository at http://bib.umassmed.edu/influenza/.

## Results and Discussion

### Effects of skewed offspring distributions on variation within populations

To underscore the importance of properly accounting for skewed offspring distributions when inferring selection from population genetic data, we briefly illlustrate the effects of sweepstakes reproduction on two population genetic summary statistics. Under a model of sweepstakes reproduction where the variable *Ψ* describes the proportion of individuals in a generation that are the offspring of a single individual in the previous generation, we summarize in Fig. 1 the SFS and Tajima’s *D*, averaged over 100 replicate populations of size *N* = 1000 under a broad range of *Ψ*.

The primary points of note are that under equilibrium neutrality non-zero values of *Ψ* skew the SFS toward an excess of singletons and high-frequency variants, and that Tajima’s *D* is negatively correlated with *Ψ*. The reader may note that Tajima’s *D* is slightly negative for *Ψ* = 0, as should be expected given that Tajima’s *D* is a biased summary of the SFS dependent upon the recombination rate (Thornton, 2005).

Hence, it is clear that failure to account for offspring skew may result in mis-inference, as null model expectations strongly differ from those of the WF model. In the following sections, we will demonstrate that accounting for sweepstakes reproduction simply as a decrease in *N_e_* (as in WF-ABC) results in highly biased estimates of selection. However, explicitly incorporating the underlying processes of MMC events can correctly adjust for their effects and yield accurate and precise estimates of *s* from time-series data.

### Estimation of Ψ with MMC-ABC

In Step 1 of MMC-ABC, the trajectories of all sites included in the data are used to estimate *N_e_* using the unbiased estimator of Jorde and Ryman (2007). In the case where the census size or harmonic mean of the population size across all time points is known, as is often the case in experimental lineages, populations of census size *N* with sweepstakes parameter *Ψ* drawn from its prior and mutational frequencies matching those at the first time point of the data are simulated for the same number of generations as the original data. The best 1% of simulations are retained to generate a posterior for *Ψ*.

MMC-ABC is able to accurately infer *Ψ* over a broad parameter space. Fig. 2 shows the mean of the posterior distribution of *Ψ* averaged over 1000 replicate populations each at *Ψ* ∈ {0, 0.01,…0.25} in the case where the correct value of *N* is specified. These illustrative parameter values were chosen to match general features of common viral experimental evolution studies (e.g., Foll et al., 2014a; Bank et al., 2016; Ormond et al., 2017)

**Figure 2.**
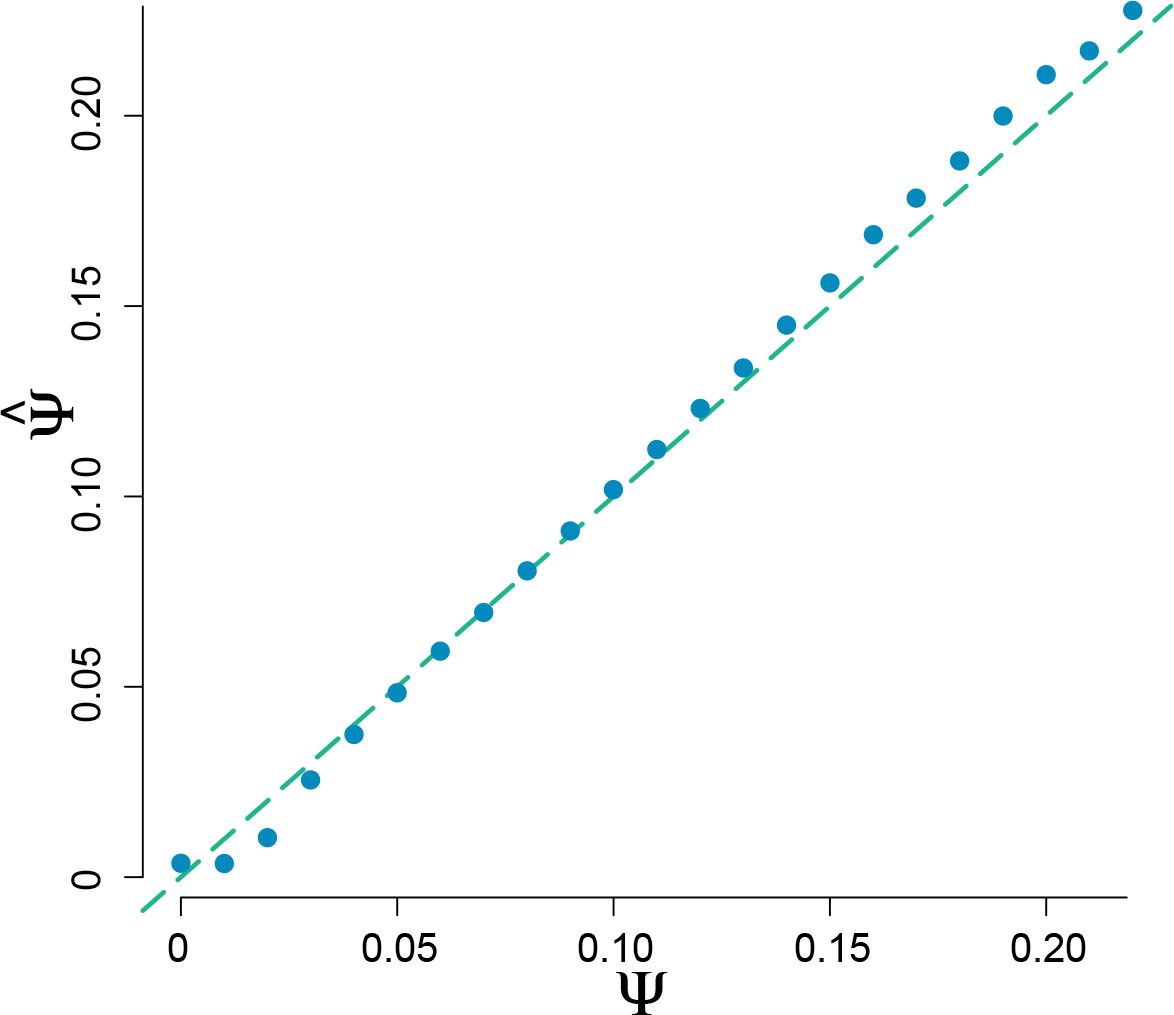
Estimation of *Ψ* by MMC-ABC, with estimates averaged over 1000 replicate populations of size *N* = 1000, with an average of 300 polymorphic sites per population tracked at 20 time points over 200 generations with a minimum of 8 informative time points, with the correct value of *N* specified. True values of *Ψ* are indicated by the dashed line. Thus, MMC-ABC accurately estimates the value of *Ψ* from time-series data when the true value of *N* is known.

Although in cases of experimental evolution precise measurements of *N* may be available to inform the prior used in Step 1 of MMC-ABC, knowledge of the size of the population in question may not be available. Therefore, we determined the power of MMC-ABC to accurately estimate *Ψ* in the absence of knowledge about the true value of *N*. In this case, both *N* and *Ψ* are drawn from priors, and MMC-ABC generates a joint posterior for the two parameters. We found that MMC-ABC is a good estimator of *Ψ* even when a large, uniform prior is used (Fig. 3). MMC-ABC likewise performs well in the case where a single, incorrect value of *N* is specified, particularly for high values of *Ψ*, at which 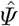 converges at the true value due to the non-linear relationship between *Ψ* and *N_e_* (Fig. 3).

**Figure 3.**
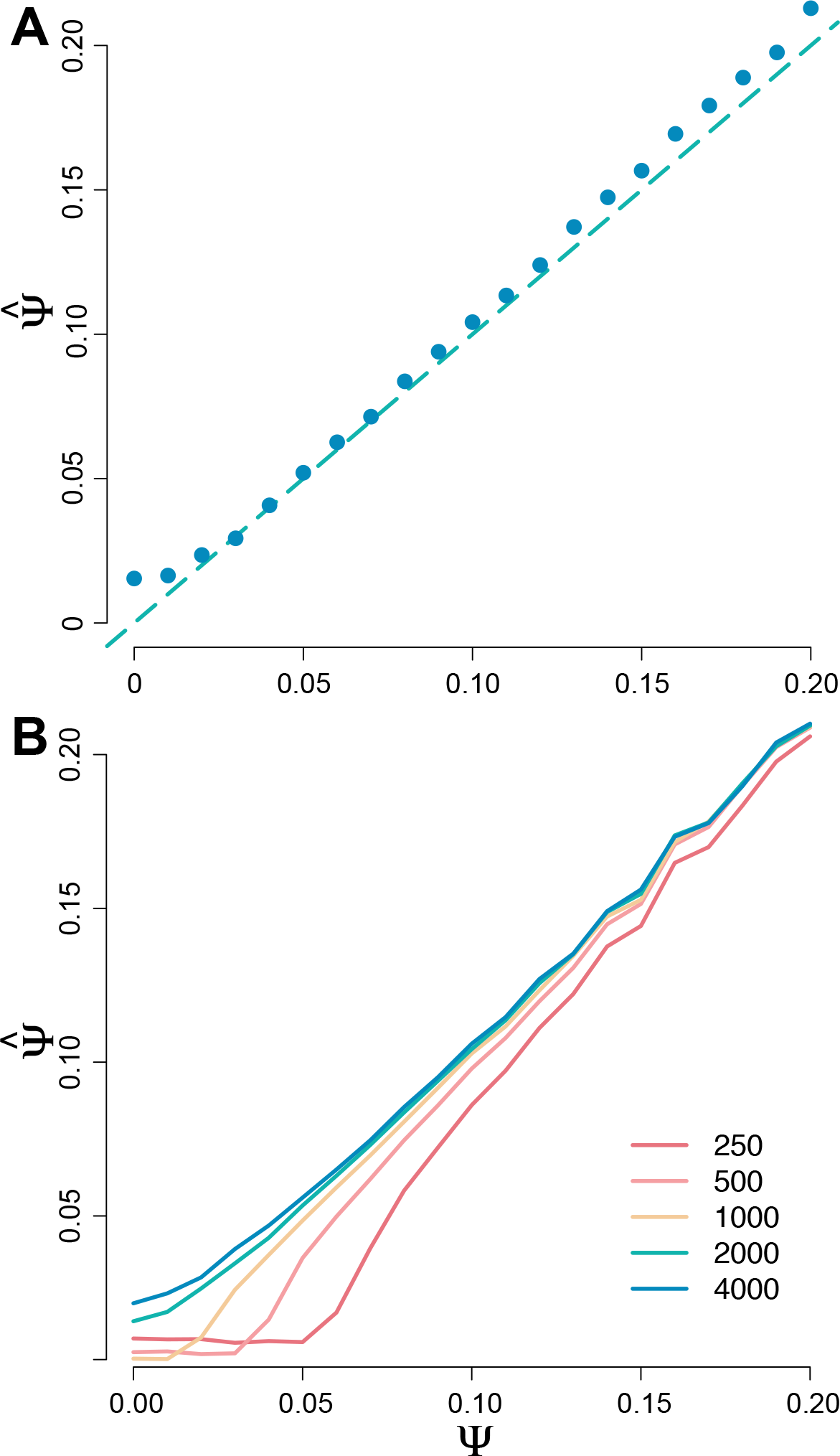
**A**: Estimation of *Ψ* by MMC-ABC, with estimates averaged over 100 replicate populations of size *N* = 1000 with values of *Ψ* and *N* drawn from priors *~U* [0, 0.3] and *~U* [250, 4000], demonstrating the robustness of MMC-ABC to mis-specification of census size. True values of *Ψ* are indicated by the dashed line. **B:** Estimation of *Ψ* by MMC-ABC, with estimates averaged over 100 replicate populations of size *N* = 1000 with either the correct value of *N* or an incorrect value of *N* (*N* ∈ {250, 500, 1000, 2000, 4000}) specified, demonstrating the non-linear relationship between *N* and *N_e_* under the Ψ- coalescent, with mis-specification of *N* having little effect on the accurate estimation of *Ψ* when *Ψ* is large.

We assessed the performance of MMC-ABC over a range of data types, including cases with 5, 11, or 21 time points over a span of 100 generations, as well as for sample sizes of 25, 100, and 250 for populations of *N* = 1000 at *Ψ* ∈ {0, 0.05, 0.1, 0.15, 0.2} (Figs. S1; S2). As expected, the estimation of *Ψ* improves with larger sample sizes and more densely sampled time points. However, MMC-ABC remains a good estimator of *Ψ* even with as few as five time points or a sample size of 25.

### Estimation of site-specific selection coefficients

In the second step of MMC-ABC, the posterior distributions of *N* and *Ψ* obtained in Step 1 are used to simulate 10,000 trajectories at each site *x_i_* (*i* = 1,…, *L*) with the alleles introduced in the population at the initial frequency *x*_*i*,1_ provided in the data. The best 1% of simulations are retained to generate a posterior for *s*. Foll et al. (2014b) previously demonstrated WF-ABC to be a good estimator of genome-wide *N_e_* and site-specific selection coefficients in populations well-described by the Kingman coalescent. The performance of WF-ABC matched or exceeded that of other similar methods. Therefore, we restrict our comparison of the performance of MMC-ABC to that of WF-ABC. For a detailed comparison of the performance of WF-ABC with that of other methods, including those of Bollback et al. (2008), Malaspinas et al. (2012), and Mathieson and McVean (2013), see the results of Foll et al. (2014b).

To compare the ability of MMC-ABC and WF-ABC to infer site-specific selection coefficients, we estimated *s* for 1000 trajectories simulated under the Ψ-coalescent with *N* = 1000 and *Ψ* = 0.1 for *s* ∈ {0, 0.1, 0.2, 0.3, 0.4}. All allele trajectories began from a minor allele frequency of 10%. Because the summary statistics used by MMC-ABC and WF-ABC assume that the majority of sites are neutral, we provided true values of *N* and *Ψ* to MMC-ABC and of *N* to WF-ABC in this initial comparison. As shown in Fig. 4, MMC-ABC is very accurate at estimating *s* under recurrent and strong sweepstakes reproduction, while WF-ABC consistently overestimates selection coefficients for positively selected sites and underestimates *s* for neutral sites. The same is true for small values of *s* ∈ {0, 0.005, 0.01, 0.015, 0.02}.

**Figure 4.**
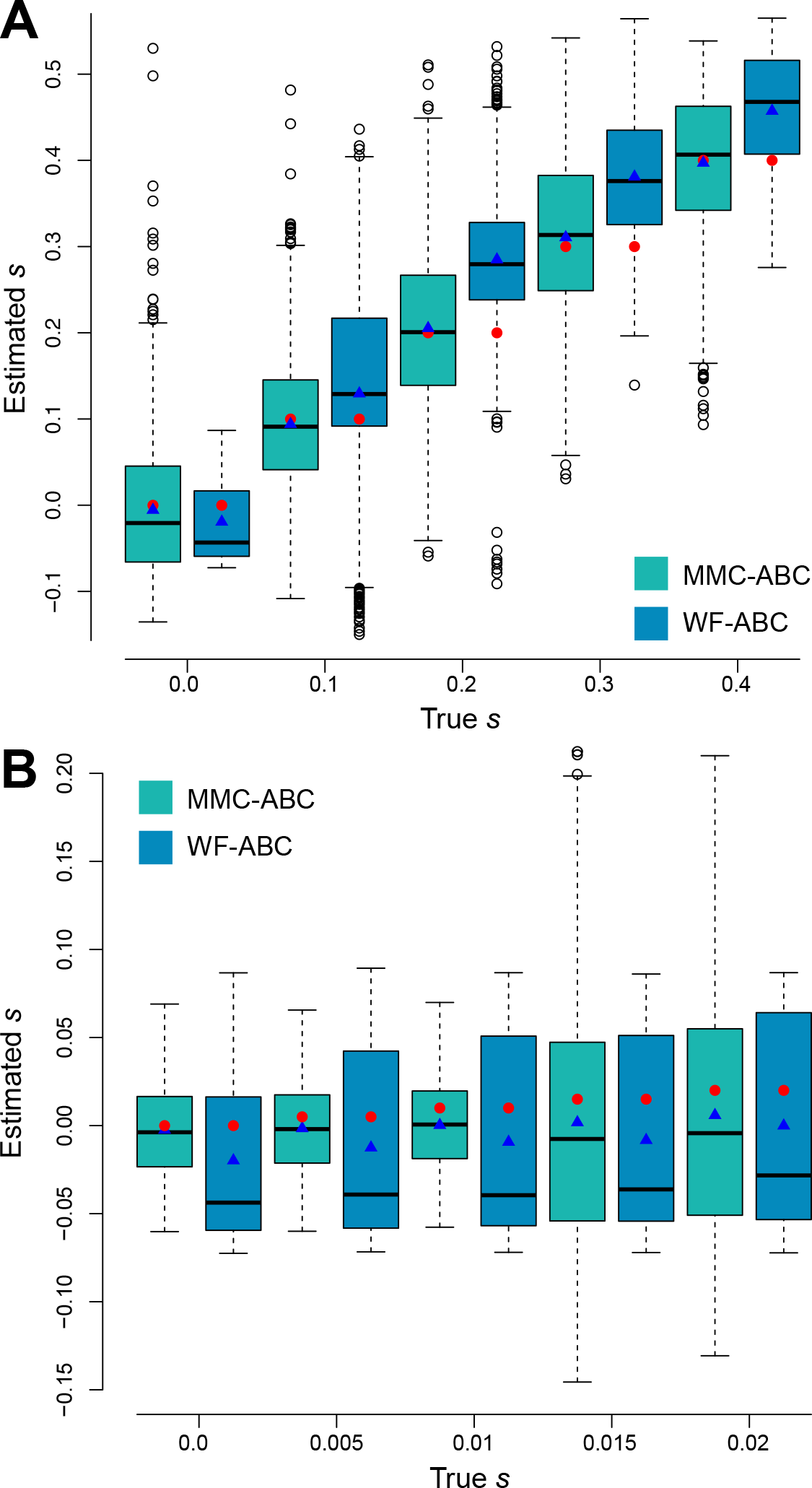
**A:** Estimation of *s* by MMC-ABC and WF-ABC for 1000 sites under selection for *s* ∈ {0, 0.1, 0.2, 0.3, 0.4} with the true values of *N* = 1000 and *Ψ* = 0.1 provided to MMC-ABC and the true value of *N* provided to WF-ABC. Results presented in a standard box plot with the box as the first, second and the third quartiles, and the whiskers as the lowest and highest datum within the 1.5 interquartile range of the lower and upper quartiles, respectively. Red circles indicate the true value of *s*, and blue triangles indicate the sample mean. **B:** Estimation for *s* ∈ {0, 0.005, 0.01, 0.015, 0.02} with the same conditions as above. WF-ABC tends to underestimate *s* for neutral alleles and overestimate *s* under strong positive selection under sweepstakes reproduction.

Estimating *s* for single trajectories of mutations covering a wider range of true selection coefficients from −0.1 to 0.4, it is evident that MMC-ABC is not only a good estimator under sweepstakes reproduction of selection for sites under positive selection and neutrality, but is also accurate for sites under negative selection. WF-ABC, however, in addition to having a strong bias toward overestimation of *s* for sites under positive selection, is negatively biased for neutral and negatively selected sites (Fig. 5). Inference under the Kingman for organisms that violate the assumption of small variance in progeny distributions is thus prone to serious over-or under-estimation of selection coefficients, while correctly accounting for reproductive skew produces accurate estimates of selective strength. This mis-inference under the WF model results from the acceleration of transit times under sweepstakes reproduction, which are interpreted by WF-ABC as an amplification of positive or negative selection.

**Figure 5.**
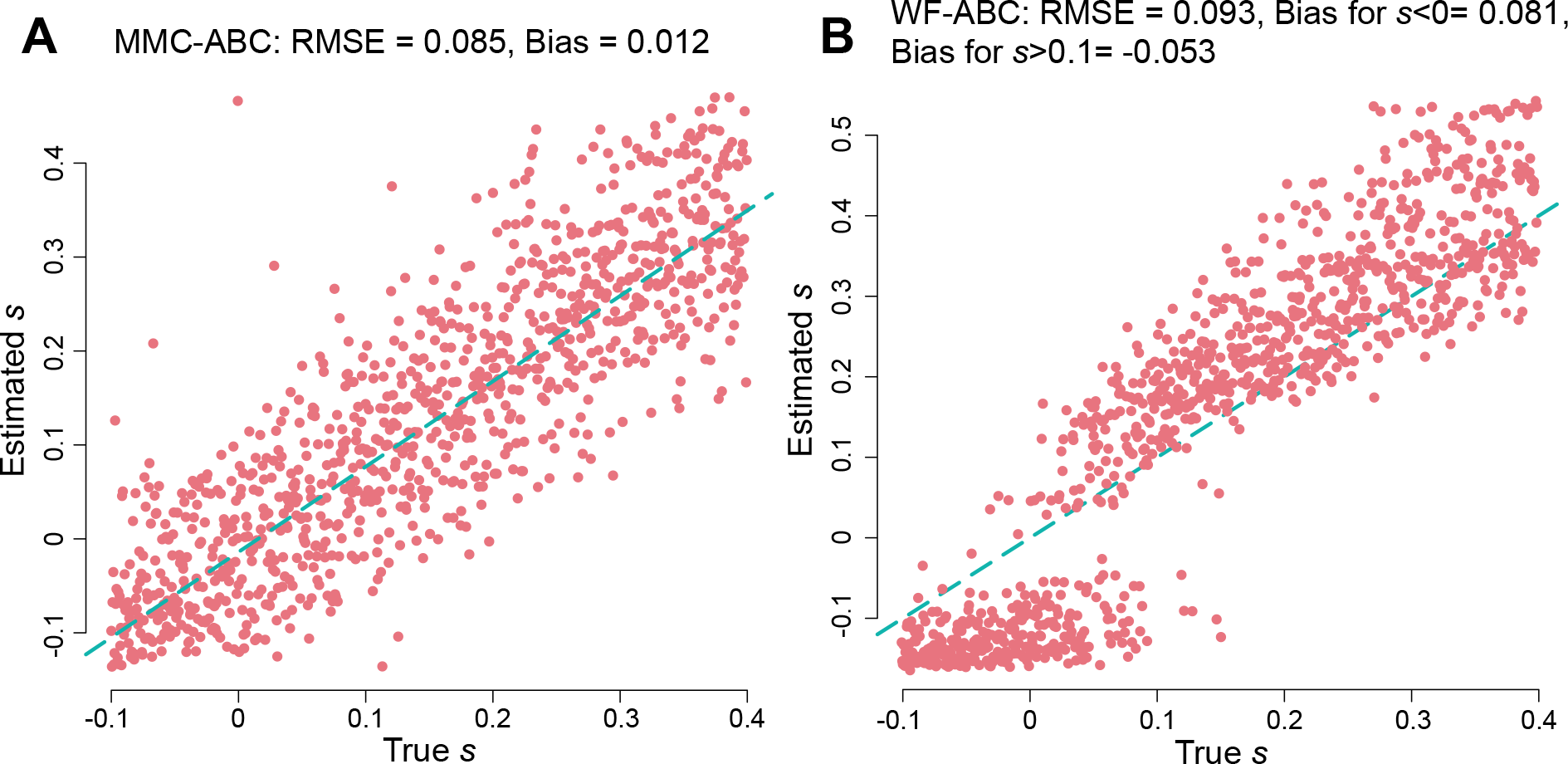
Estimation of *s* by MMC-ABC (A) and WF-ABC (B) for 1000 sites under selection with *s* ranging from −0.1 to 0.4, estimated over 10,000 simulated values of *s* for each site, and with the correct values of *Ψ* = 0.1 and *N* provided to MMC-ABC. MMC-ABC is a relatively unbiased estimator of *s* under offspring skew, while WF-ABC strongly overestimates *s* for positively selected sites and underestimates *s* for neutral and negatively selected sites.

As with the estimation of *Ψ*, we assessed the performance of Part 2 of MMC-ABC over a range of data types, including cases with 5, 11, or 21 time points over a span of 100 generations, as well as for sample sizes of 25, 100, and 250 for populations of *N* = 1000 at *s* ∈ {0, 0.1, 0.2, 0.3, 0.4} (Figs. S3; S4). Again, as expected, the estimation of *s* improves with larger sample sizes and more densely sampled time points. However, MMC-ABC is a reasonably good estimator of *s* even with as few as five time points or a sample size of 25.

### Joint estimation of *N*, Ψ, and *s*

We simulated trajectories for 9500 neutral loci and 500 selected loci for which *s* = 0.1, under conditions in which *N* = 1000 and *Ψ* = 0.1. MMC-ABC estimated firstly *N* and *Ψ* over priors of *~U* [250, 2000] and *~U* [0, 0.3], respectively, and then estimated *s* for each site with values of *N* and *Ψ* drawn from the joint posterior (Fig. 6). The estimated value of 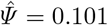, the mean estimated value of *s* for neutral sites was 0.008 and for positively selected sites was 0.924, highlighting the ability of MMC-ABC to jointly estimate the magnitude of skewed offspring distributions and site-specific selection coefficients with accuracy, even when a relatively large proportion (5% in this case) of sites are under strong positive selection.

**Figure 6.**
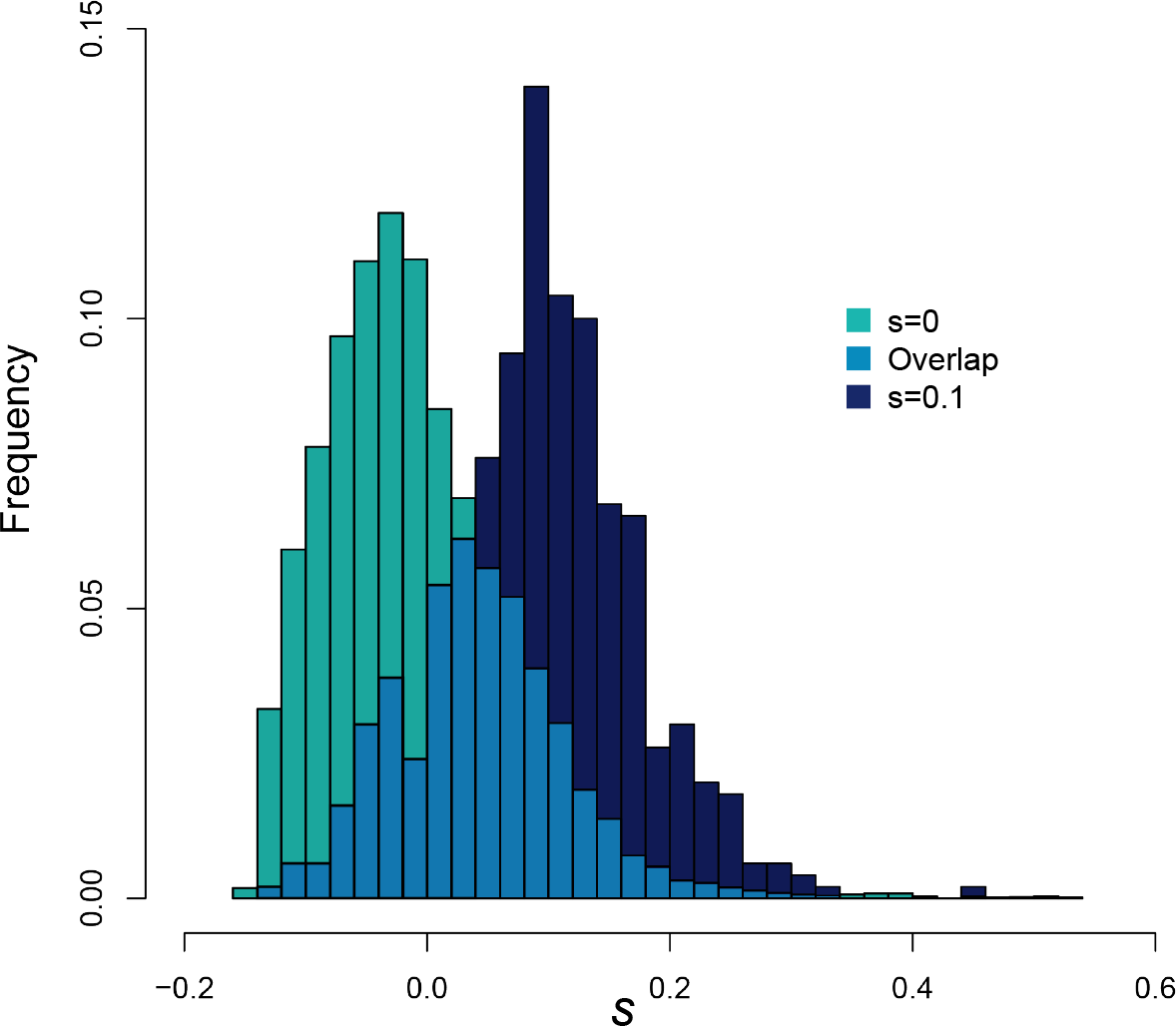
Estimation of *s* by MMC-ABC for 9500 hundred neutral sites and 500 sites for which *s* = 0.1 with *N* = 1000 and *Ψ* = 0.1 with *N* estimated over a uniform prior *~U*[250, 2000], *Ψ* estimated from the prior *~U*[0, 0.3] and *s* estimated over *~U* [−0.2, 0.6]. Note that we display the relative frequencies for estimated values of *s* for each class of mutation, for which there were unequal numbers of total sites. These results demonstrate the ability of MMC-ABC to jointly and accurately estimate *N*, *Ψ*, and *s* from genomic data, even when a large number of sites are under positive selection.

This result is notable, given that both recurrent positive selection and skewed progeny distributions can result in coalescent trees dominated by multiple-mergers (Durrett and Schweinsberg, 2004, 2005). Different features of the data—resulting from the localized effects of selection and the genome-wide effects of sweepstakes reproduction—allow us to disentangle the MMC behavior of neutral offspring skew from that of non-neutral offspring skew generated by positive selection.

### Application to data from *influenza A*

We applied MMC-ABC to time-series data from the experimental evolution of influenza A. These data were collected under standard culture conditions and during a period of exposure to exponentially increasing concentrations of the drug oseltamivir (Foll et al., 2014a).

The data consist of time-sampled minor allele frequencies for two control lineages and two drug-selected lineages. Using WF-ABC, Foll et al. (2014a) previously estimated the effective population sizes of the control and selected populations to be 176 and 226, respectively, with values of *N_e_* derived from the harmonic means of the population sizes during passaging being 737 and 696, respectively. They hypothesized that the discrepancies in measurements of *N_e_* were likely due to the large variance in viral burst sizes, yielding skewed offspring distributions. These experimentally evolved populations therefore are well-suited to the application of MMC-ABC.

We first obtained estimates of *Ψ* for each population, using the harmonic population size means as a prior for *N*. We then obtained posterior distributions of *s* for all mutations segregating in at least two time points and with a minimum frequency of 2.5% for at least one time point. We define Bayseian ‘*p*-values’ for *s* as *P* (*s* < 0|*x*) and consider a trajectory to be ‘significant at level *p*’ if its equal-tailed 100(1 − *p*)% posterior interval excludes zero (Beaumont and Balding, 2004).

The mean posterior estimate of *Ψ* for the two control lines was 0.067, and the mean value of *Ψ* across both drug-treatment lines was 0.084. MMC-ABC recovered two of the same six control line mutations and seven of the 15 mutations from the drug selection lines identified by (Foll et al., 2014a) as being beneficial at the level *p* = 0.01 (Table 1, summarizing all eight mutations significant under MMC-ABC and eight of the 20 significant under WF-ABC, sites of known drug-resistance mutations shown in bold font). The mutations of significant beneficial effect under WF-ABC had an average effect of *s* = 0.1 for control line mutations and *s* = 0.13 for drug-selection mutations. The same sets of mutations (including those that did not achieve significance under MMC-ABC) had average effects of *s* = 0.11 and *s* = 0.17 as estimated by MMC-ABC.

**Table 1.**
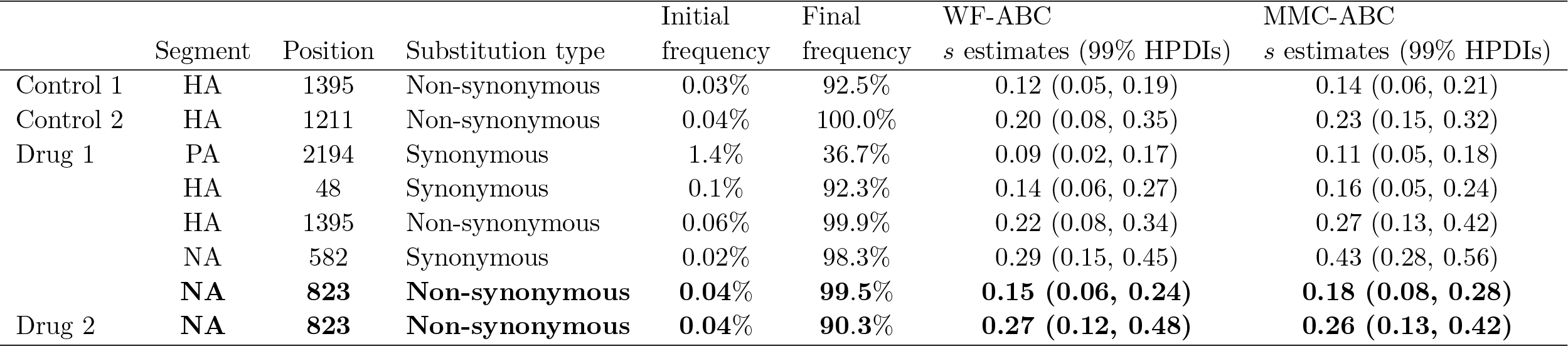
Mutations significantly beneficial under MMC-ABC from experimentally evolved populations of influenza A virus

The beneficial mutations identified in the control lines are likely adaptations to the MDCK cells used in serial passaging. One mutation, which rose to high frequency in the first control and drug lines at nucleotide position 1395 of the hemagglutinin segment, has been widely observed across influenza strains and is a common adaptation to tissue culture (Daniels et al., 1985; Reed et al., 2009; Foll et al., 2014a). Another mutation, which reached high frequency in the second control line, has likewise been associated with adaptation to culture conditions (Lin et al., 1997; Ilyushina et al., 2007). Notably, the mutation at position 823 of the neuraminidase segment (identified as H275Y under the N2 numbering system) achieved high frequency in both drug lineages and is a well-documented resistance mutation for oseltamivir (Sha and Luo, 1997; Arzt et al., 2001; Collins et al., 2008).

Six of the eight synonymous mutations found to be significantly beneficial by WF-ABC were not significantly beneficial under MMC-ABC. By estimating an appropriate neutral null model under the Ψ-coalescent, we reduced the list of candidate resistance mutations, thus likely minimizing the rate of false positives and excluding many hitchhiking mutations (as the synonymous sites are likely to be). Several experimentally validated mutations known to improve either infectivity in tissue culture or resistance to oseltamivir were retained under MMC-ABC, as were a handful of other potential candidate resistance mutations.

## Conclusions

The revolution in sequencing technology has increased the availability of time-series polymorphism data by orders of magnitude, but the utility of such data relies upon the derivation and development of appropriate inference methodologies. The neutral biology of large swaths of the tree of life renders the most common class of inference based on the Kingman coalescent of questionable use. We have demonstrated here that performing inference under the assumptions of the Wright-Fisher model and the Kingman coalescent leads to an incorrect understanding of both population size and selection coefficients in such organisms. Matuszewski et al. (2018) have also shown the same to be true for the demographic history of the population. Fortunately, the theoretical details are in place to develop similar inference of demography and selection under biologically appropriate alternative coalescent models (Wakeley, 2013).

We have shown that MMC-ABC is able to jointly estimate *N*, *Ψ*, and site-specific selection coefficients accurately, even under high levels of reproductive skew and with an unknown population size. Notably, we are able to distinguish selection-induced offspring skew from skew originating from the neutral reproductive biology of populations, largely due to the genome-wide scale of MMC events relative to the localized effects of selection. We are also able to differentiate drift-induced effects imposed by small population sizes from those induced by sweepstakes reproduction events.

Very little is known regarding the extent of progeny skew across groups of viruses, bacteria, and plants, or the extent of skew artificially induced by domestication and cultivation. However, this work demonstrates that, at least with time-sampled allele frequency data, such inference is now possible, and moreover will allow for the construction of much more accurate neutral null models in these organisms, which will greatly reduce false-positive rates in scans for selection, provide a more accurate picture of demographic history, and reveal previously hidden details regarding variance in offspring number.

## Acknowledgements

This work was funded by grants from the European Research Council, the Swiss National Science Foundation, and the U.S. Department of Defense to JDJ. We thank Stefan Laurent for helpful discussion.

